# Structural plasticity of CENP-A regulated by H4 influences cellular levels and kinetochore assembly

**DOI:** 10.1101/282863

**Authors:** Nikita Malik, Sarath Chandra Dantu, Mamta Kombrabail, Santanu Kumar Ghosh, Guruswamy Krishnamoorthy, Ashutosh Kumar

## Abstract

The Histone variant CENP-A^Cse4^ is a core component of the specialized nucleosome at the centromere in budding yeast. The level of Cse4 in cells is tightly regulated, primarily by ubiquitin-mediated proteolysis. However, the structural transitions in Cse4 that regulate centromere localization and interaction with regulatory components are poorly understood. Using time resolved fluorescence, NMR and molecular dynamics we show for the first time that soluble Cse4 can exist in a ‘closed’ conformation, inaccessible to various regulatory components. We further determined that binding of its obligate partner H4, alters the inter-domain interaction within Cse4, ensuring an ‘open’ state that will lend itself to proteolysis. This dynamic model allows kinetochore formation only in presence of H4, as the N-terminus, which is required for interaction with centromeric components will be unavailable in absence of H4. The specific requirement of H4 binding for the conformational regulation of Cse4 suggests a unique structure-based regulatory mechanism for Cse4 localization and prevention of premature kinetochore assembly.

## Introduction

The basis of successful cell division is the faithful segregation of sister chromatids during mitosis and meiosis, a process driven by the formation of the kinetochore complex on the centromere. Centromeres in most eukaryotes are identified by the formation of specialized nucleosome(s) where the Histone 3 (H3) is replaced by a unique variant CENP-A (Centromeric protein-A) [1]. In budding yeast this variant, known as Cse4 [2] forms the specialized nucleosome [3,4] at a single centromere that mediates the segregation of chromosomes [5]. The localization of Cse4 at the centromere, and its level in the cells has to be tightly regulated, as altered localization and expression of this protein is known to cause genetic instability [6]. Two distinct pools of Cse4 are present in the cell; the core Cse4 at the centromere that mediates kinetochore formation and a pericentromeric reservoir that provides Cse4 molecules in case of eviction of core molecules from the centromere [7]. How these populations are maintained in the cells and how the peri-centromeric Cse4 is localized at the centromere, is not clearly understood. Ubiquitin mediated proteolysis is one of the key mechanisms that are known to regulate Cse4 levels and maintaining kinetochore function [8]. Psh1 was identified as the E3 ligase that specifically recognizes the CENP-A Targeting Domain (CATD) in Cse4 and prevents its misincorporation in the chromatin [9,10]. Recently, an evolutionarily conserved protein Pat1 was identified that protects Cse4 from Psh1 mediated degradation and is thought to maintain the population of pericentric Cse4 molecules at the kinetochore [11]. Interestingly, in the above studies, Cse4 is not completely stabilized when Psh1 is deleted, and a lysine free mutant of Cse4 still gets degraded in the cell. Some other proteins like Doa1/Ufd3 and Rcy1 are also implicated to be essential for Cse4 proteolysis [12,13]. However, none of the mechanisms have shown complete regulation of the Cse4 levels. Thus it is possible that there exist other, perhaps ubiquitin-independent mechanisms in the cell that are regulating the Cse4 levels.

We asked if the conformation of the soluble Cse4 molecule itself could be a determining factor in mediating its interaction with regulators and with the kinetochore machinery since, the structural basis of these interactions is not clear. The C-terminus of Cse4 is essential for centromere targeting [14,15] and has thus been the focus of many studies. The structure of the Histone fold domain (HFD) within the C-terminus of Cse4 in complex with Histone 4 (H4), and the chaperone Scm3 has been elucidated [16–18], but very little is known about the conformation of the N-terminus of Cse4. Cse4 has a unique, long N-terminus, which harbors the Essential N-terminal Domain (END) (residues 28-60), required for interaction with other kinetochore proteins [19]. We hypothesized that the 129 a.a N-terminal domain (NTD) could have a more specialized function than other histone tails and that a longer tail could have a structural role in regulating the association of Cse4 with other proteins. Using time resolved fluorescence, NMR and molecular dynamics we found that in native state the NTD interacts with the C-terminus domain (CTD). Such an interaction constitutes what we call a ‘closed’ conformation of the Cse4 monomer and has not, to our knowledge, been reported for any histone so far. Such a conformation may hinder the association of regulators with the CENP-A Targeting Domain and/or that of kinetochore proteins with the NTD, thereby negating the untimely mis-targeting of Cse4 within the nucleosome and to other ectopic loci. In addition, we investigated whether the ‘closed’ conformation of Cse4 was altered in the presence of H4, its obligate partner in the specialized nucleosome [4,20]. We observed that H4 binding indeed curtailed the interaction between the NTD and CTD domains of Cse4. The interaction with H4 allows the transition to an open conformation permitting the interaction of Cse4 with kinetochore proteins to assemble a kinetochore and allowing the degradation machinery to access the CTD in case of any inadvertent mis-localization. Our results thus suggest a novel structure-based mechanism based on the conformational flexibility of the N-terminus, to regulate the levels of Cse4 in the cell and to resist premature kinetochore assembly.

## Results

### N-terminus of Cse4 is conformationaly restricted

In order to understand the effect of other centromeric components on the NTD of Cse4 (1-129) it is vital to understand its behavior in the context of the Cse4 monomer. Cse4 contains two Trp residues in its sequence, W7 (in the NTD) which according to the modeled structure [21], is in a disordered region, whereas W178 (part of the HFD) is surrounded by side chains of neighboring residues; hence a clear difference in the fluorescence parameters of the residues is expected in the native state. The two Trp residues of Cse4 were used as intrinsic fluorescent probes by creating the single mutants W7L and W178L. In the Cse4 W178L mutant, the fluorescence signal will derive solely from the N-terminus (W7) whereas the Cse4 W7L mutant will report the behavior of the C-terminus (W178) (Fig 1a). The secondary structure of both mutants (W7L and W178L) was comparable to the wild type Cse4 (Fig S1). Surprisingly, the fluorescence lifetime values for the Trp residues at the NTD and the CTD did not show significant difference (Fig 1b, Table 1), indicating that the two residues experience similar microenvironments. As an additional measure, we calculated the solvent accessibility of the Trp residues where the N-terminus should show a higher value of the bimolecular rate constant for quenching (*k*_*q*_) [22] compared to the C-terminus if W7 is solvent exposed. However, the difference in rate constants between W178L (3.8×10^9^ M^-1^ s^-1^) and W7L (2.9×10^9^ M^-1^ s^-1^) was not significant (Fig 1b), but the *k*_*q*_ for the respective Trp residues was higher in the denatured state (Fig S1). Thus, it can be concluded that W7 is not completely solvent exposed, implying that the NTD is conformationaly restricted in the Cse4 monomer.

**Table 1:** Fluorescence intensity decay parameters for N-terminus (W178L) and C-terminus (W7L) in native state and in presence of Denaturant (D) (8 M urea)

| | Mutant | $\tau_1$ (ns) ($\alpha_1$) | $\tau_2$ (ns) ($\alpha_2$) | $\tau_3$ (ns) ($\alpha_3$) | $\tau_m$ (ns) |
| --- | --- | --- | --- | --- | --- |
| N-terminus | W178L | 0.5(0.37) | 1.7(0.48) | 4.8(0.15) | 1.7±0.1 |
| | W178L+D | 0.56(0.26) | 1.58(0.52) | 4.4(0.22) | 1.9±0.08 |
| C-terminus | W7L | 0.5(0.37) | 1.7(0.5) | 4.8(0.13) | 1.7±0.02 |
| | W7L+D | 0.67(0.31) | 1.8(0.51) | 4.9(0.18) | 2.0±0.28 |

**Fig 1.**
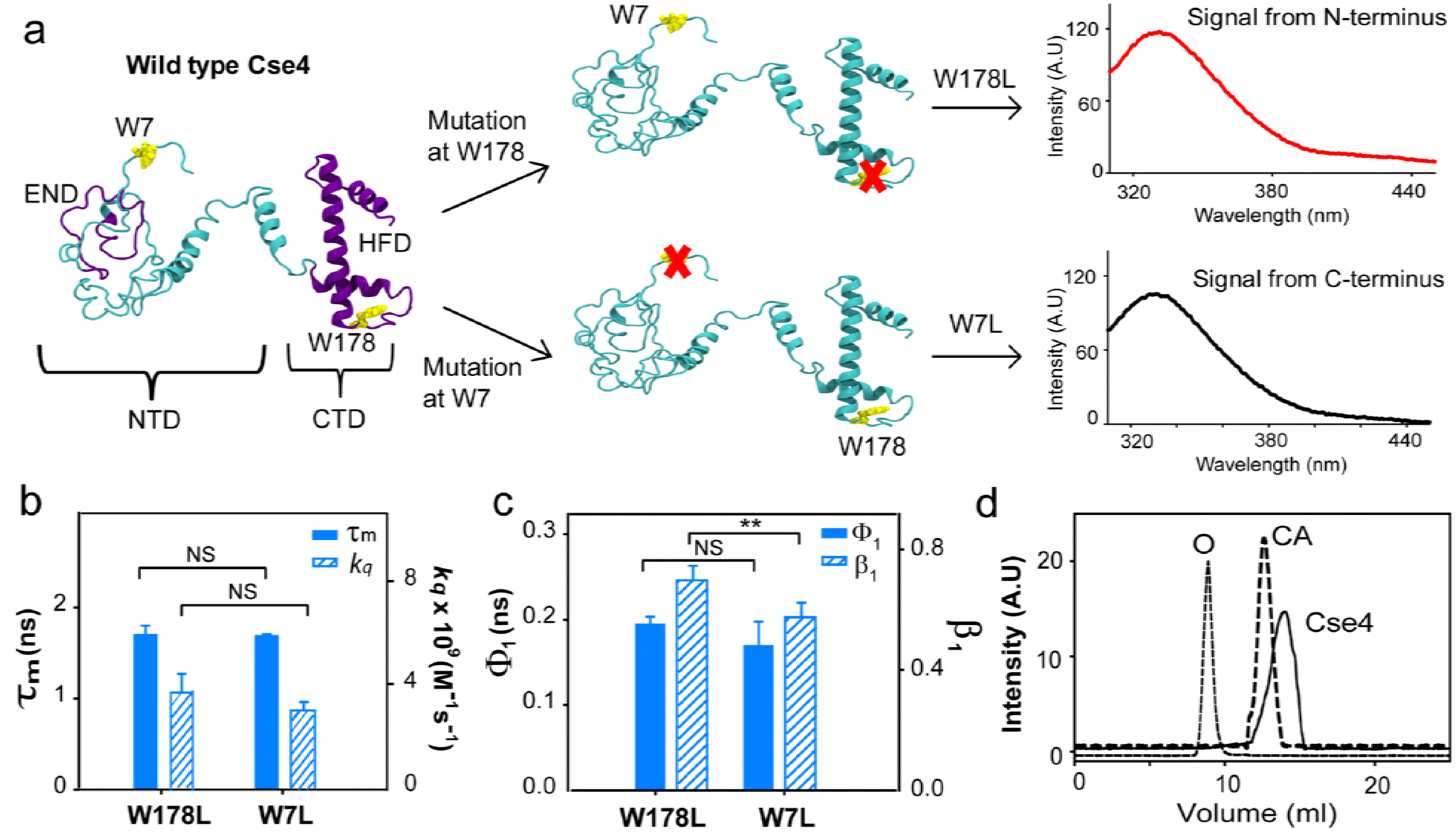
CENP-A^Cse4^ N-terminus tail is restricted. **a.** Strategy for fluorescence assay to create single Trp mutants in Cse4 to study the two domains individually, the END and HFD are highlighted in violet, NTD and CTD correspond to residues 1-129 and 130-229 respectively, note that according to a proposed model, W7 is in a disordered region implying that it is expected to have more conformational freedom than W178; **b.** Comparing fluorescence lifetimes (filled bars) and solvent accessibility (striped bars) of the two domains; **c**. Conformational flexibility of the two Trp residues in the native state; filled bars represent φ_1_ and the striped bars represent β_1_; **d.** gel filtration profile of folded Cse4 protein (solid line), the dotted line represent the protein markers (O-ovalbumin 44 kDa and CA-carbonic anhydrase 29 kDa). The statistical significance was calculated by one-way analysis of variance: *, p <0.05; **, p <0.01; NS (not significant), p >0.05; error bars represent standard deviation (SD).

To gain further insight, the conformational flexibility of the Trp residues was probed using time resolved fluorescence anisotropy. The local motion of the Trp in the protein (shorter correlation time φ_1_) and the global tumbling motion of the entire protein (longer correlation time φ_2_) contribute to the fluorescence anisotropy decay [23]. φ_1_ offers information about the site-specific conformational flexibility of the protein; higher values for φ1 and/or smaller value of its amplitude β_1_ indicate reduced flexibility. As per the Cse4 modeled structure [21], W7 is free to rotate and W178 is restricted by neighboring residues (Fig 1a); but we observed no variation in the φ_1_. A slight change in β_1_ suggested that W7 had more freedom to rotate than W178 but it still did not equate to the difference expected between a completely restricted Trp and a free Trp (Fig 1c, Table 2). The microenvironment and conformational flexibility of the NTD indicates that the tail is not completely free as believed to be in the nucleosome structure. A single peak corresponding to the molecular weight of Cse4 in gel filtration (Fig 1d) and the long component of the anisotropy decay for both mutants (∼15 ns) (Table 2) are both consistent with a protein of molecular weight ∼30 kDa, excluding the presence of Cse4 dimers or higher order structures resulting from inter-molecular interactions that may restrict the NTD. Thus the observed behavior of W7 is either due to local structural restrictions near W7 or an interaction between the N-terminus and C-terminus in the native state. Both these states could conceivably hamper the interaction of the domains with regulators as well as other centromere components. Thus, both these possibilities were further investigated.

**Table 2:**
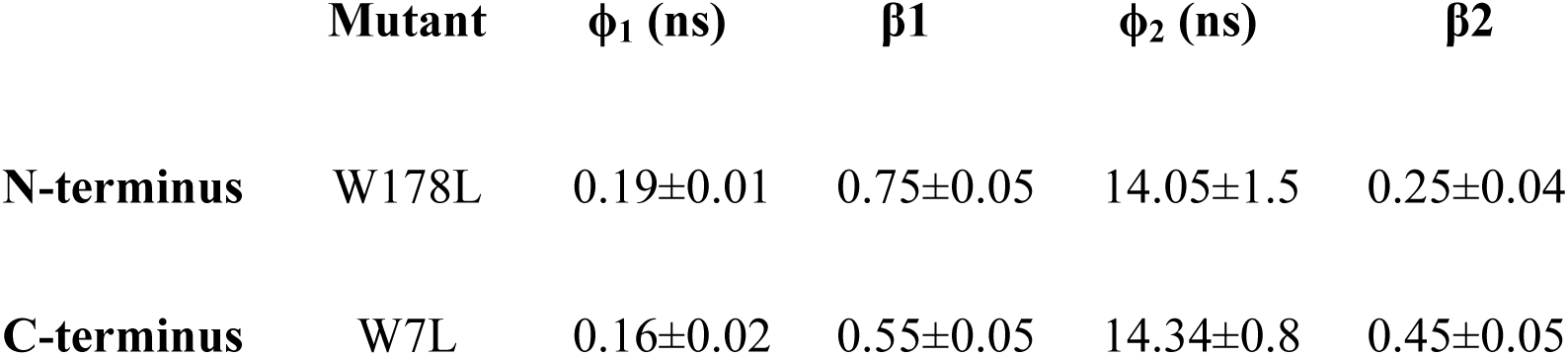
Parameters for fluorescence anisotropy decay for N-terminus (W178L) and C-terminus (W7L)

| | Mutant | $\phi_1$ (ns) | $\beta_1$ | $\phi_2$ (ns) | $\beta_2$ |
| --- | --- | --- | --- | --- | --- |
| N-terminus | W178L | 0.19±0.01 | 0.75±0.05 | 14.05±1.5 | 0.25±0.04 |
| C-terminus | W7L | 0.16±0.02 | 0.55±0.05 | 14.34±0.8 | 0.45±0.05 |

### Monomeric Cse4 exhibits inter-domain interaction

To understand whether conformational restriction of the NTD was caused by inter domain interactions between N- and C-terminus or by local structures formed at N-terminus itself, the Leu residues were changed to Ala residues in the Trp mutants mentioned above. In the event of local structural interactions within the NTD, the change in the amino acid residue at the CTD (W178L to W178A) would not affect the fluorescence parameters of Trp at the N-terminus. On the other hand, if the two domains interact, change at one terminus will have an effect on the other. It was observed that fluorescence lifetimes of the Ala mutants were significantly different from those of the Leu mutants at the C-terminus (Fig 2a, Supplementary Table S1). Similarly, there was a difference in the fluorescence anisotropy values for Trp in the two sets of mutants especially at the N-terminus (Supplementary Table S2) that showed higher conformational flexibility. The β_1_ for W178A showed a slightly smaller value than for W178L (Fig 2b). Both parameters, the microenvironment and flexibility, of Trp at one domain were affected by a change in amino acid at the other domain, suggesting that the two domains of Cse4 interact with each other. A significant difference was also observed in the *kq* values between the two mutants at the NTD, also pointing towards an interaction between the two termini (Fig 2c).

**Fig 2.**
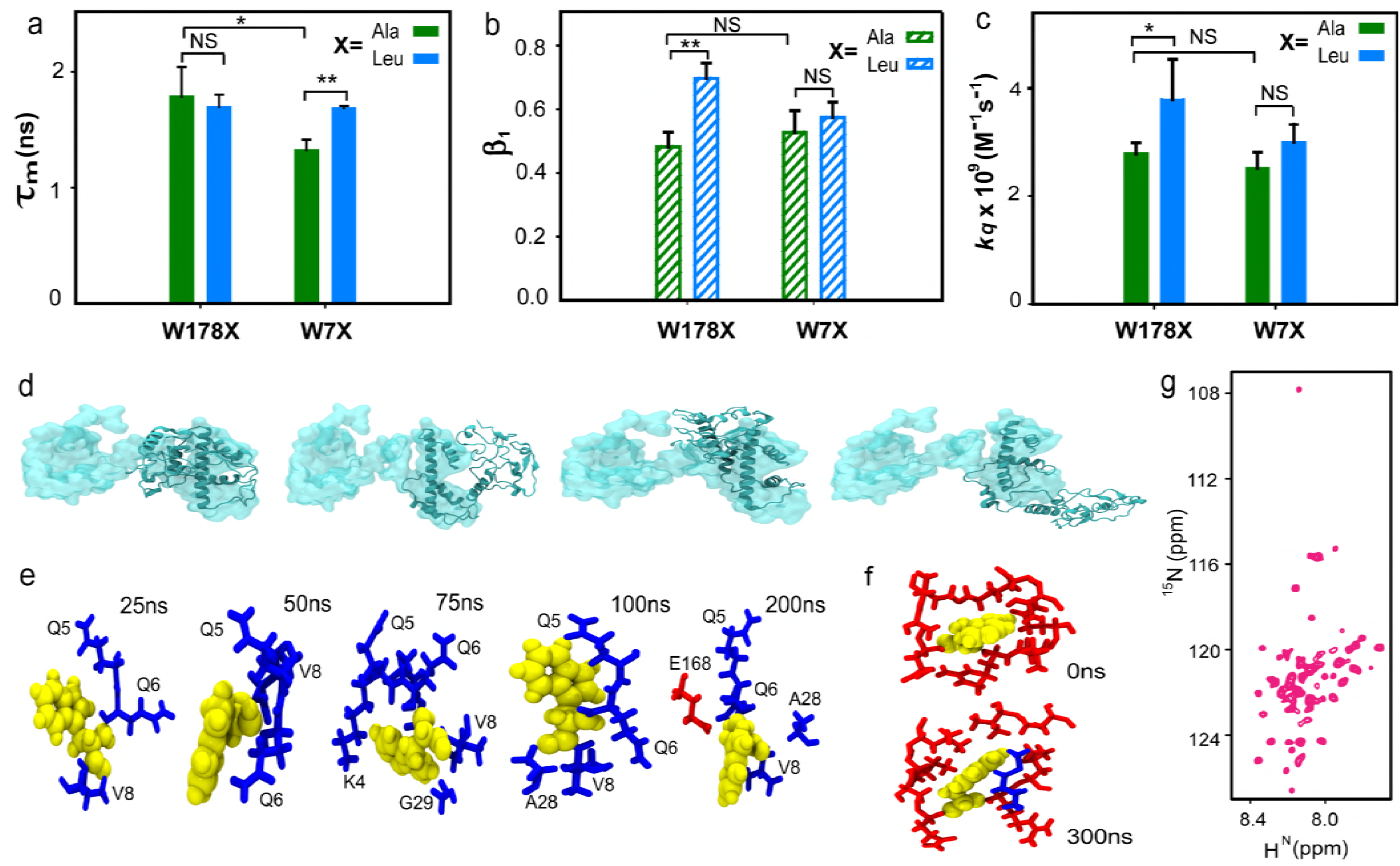
CENP-A^Cse4^ N-terminus tail interacts with C-terminus. **a**. Comparison between the Fluorescence lifetime values of the A and L mutants for the two domains; **b.** β_1_ associated with the short correlation time for each mutant; **c.** Solvent accessibility of the two domains in the A and L mutants; **d.** Overlap of the structure of Cse4 at the start (space-filled) and end (ribbon) of all four simulations, the conformation of N-terminus changes from extended to closed even though the four simulations end in different conformational basins; **e**. Change in the conformation of W7 throughout the simulation 1; **f**. Conformation of W178 at the start and end of the simulation; **g**. ^1^H-^15^N-HSQC spectrum of folded Cse4. In all graphs, X represents the mutant type with blue and green representing L and A mutants respectively. Trp residues are shown in yellow, N-terminus and C-terminus residues are shown in blue and red sticks respectively. The statistical significance was calculated by one-way analysis of variance: *, p <0.05; **, p <0.01; NS (not significant), p >0.05; error bars represent SD.

Inter-domain interaction was also observed in four independent atomistic molecular dynamics (MD) simulations of 300 ns each for monomeric Cse4 (Suplementary Movie 1). Though the four simulations ended up in different conformations at the end of 300 ns (Fig 2d, Fig S2a), the rotational flexibility of the side chain of W7 was not hindered as it interacted with different residues (Fig 2e). Even when the W7 did not directly interact with the C-terminus, the flexibility of the Trp side chain was retained (Fig S2b). This can explain the increased β_1_ observed for the N-terminus Trp. The HFD residues in vicinity (<4 Å) of W178 did not change through the simulation and in some simulations residues from the NTD were also observed in the proximity of W178 (Fig 2f). The interaction between the domains was also evident from the multiple contact points between the NTD and CTD (Fig S3a) and reduced distance between the Cα atoms of W7 and W178 (Fig S3b,c) in all the four simulations.

NMR spectroscopy was used to get an insight into residue specific dynamics and structure of the NTD. The ^15^N-HSQC spectrum of Cse4 showed fewer than the expected number of peaks (224 non-Pro peaks) (Fig 2g). However, due to inadequate sample concentration, NMR assignments could not be completed (Fig S4). We believe that the residues involved in the interaction between the two domains have broadened due to exchange, leaving only the peaks of the non-interacting residues visible in the spectrum.

### H4 binding stabilizes CENP-A^Cse4^

Since, histone tails and their cores are known to have distinctive structures and functions, we wanted to understand if the two domains of Cse4 also behave independently, as this would affect their interactions with other proteins. In order to assess whether the domains showed distinctive behavior during folding Cse4 was denatured and changes in the residue-wise secondary structure and dynamics of the protein were monitored as the denaturant (8 M urea) was diluted. Spectra were observed to shift considerably and peak broadening was seen at lower urea concentrations (Fig S5a). Resonances from the CTD started disappearing at 5 M urea, whereas those from NTD, though shifted, were still visible (Fig 3a), signifying that CTD residues undergo conformational exchange earlier than those at the N-terminus. At 4 M urea, the protein appeared to be in a molten globule state (Supplementary Fig 5a). To quantify these changes, residue-wise secondary structural propensities were calculated. The propensities at 6 M urea (Fig S5b) were similar to the protein in 8 M urea [24]. But significant changes were seen at 5 M urea concentration where patches of the NTD showed helical propensities (Fig 3b). Though the protein is not in the native conformation, these still give information about the structural rearrangements within the N-terminus residues. The results indicate that the NTD is not completely disordered, but undergoes structural transitions independent of the CTD. It should be noted that these may or may not be the native propensities of the N-terminus residues and further rearrangements are possible in the native state as observed earlier [25]. The comparison between the ^15^N R_2_ (^15^N transverse relaxation rates) of Cse4 in 8 M and 6 M urea showed a considerable increase in the values for the residues 30-51 and 190-205 (Fig S5c, Fig 3c). These residues belong to the critical END and HFD regions of Cse4 (Fig 3d), demonstrating that biologically essential regions serve as early nucleation sites for Cse4 folding. Cse4 contains five glycine residues, which are distributed throughout the 229 a.a sequence (G18, G29, G79, G196, G226). As a representative for both domains, these Gly residues were used as markers to map the dynamics of the different regions of Cse4 (Fig 3d, e). The increase in R_2_ for G196 and absence of peaks at 4 M urea reinforces the point that HFD is the early nucleation site for folding. The Gly residues at NTD (G18, G29) showed a consistent increase in R_2_ from 8 M to 4 M urea, indicating increased rigidity and/or conformational exchange that may result from its interaction with the CTD; the residue G79 does not change significantly signifying that it is not involved in the interaction. This trend was seen in their intensity and positions, where NTD Gly residues were shifted but those from CTD had broadened in 5 M urea spectra (Fig 3a, inset).

**Fig 3.**
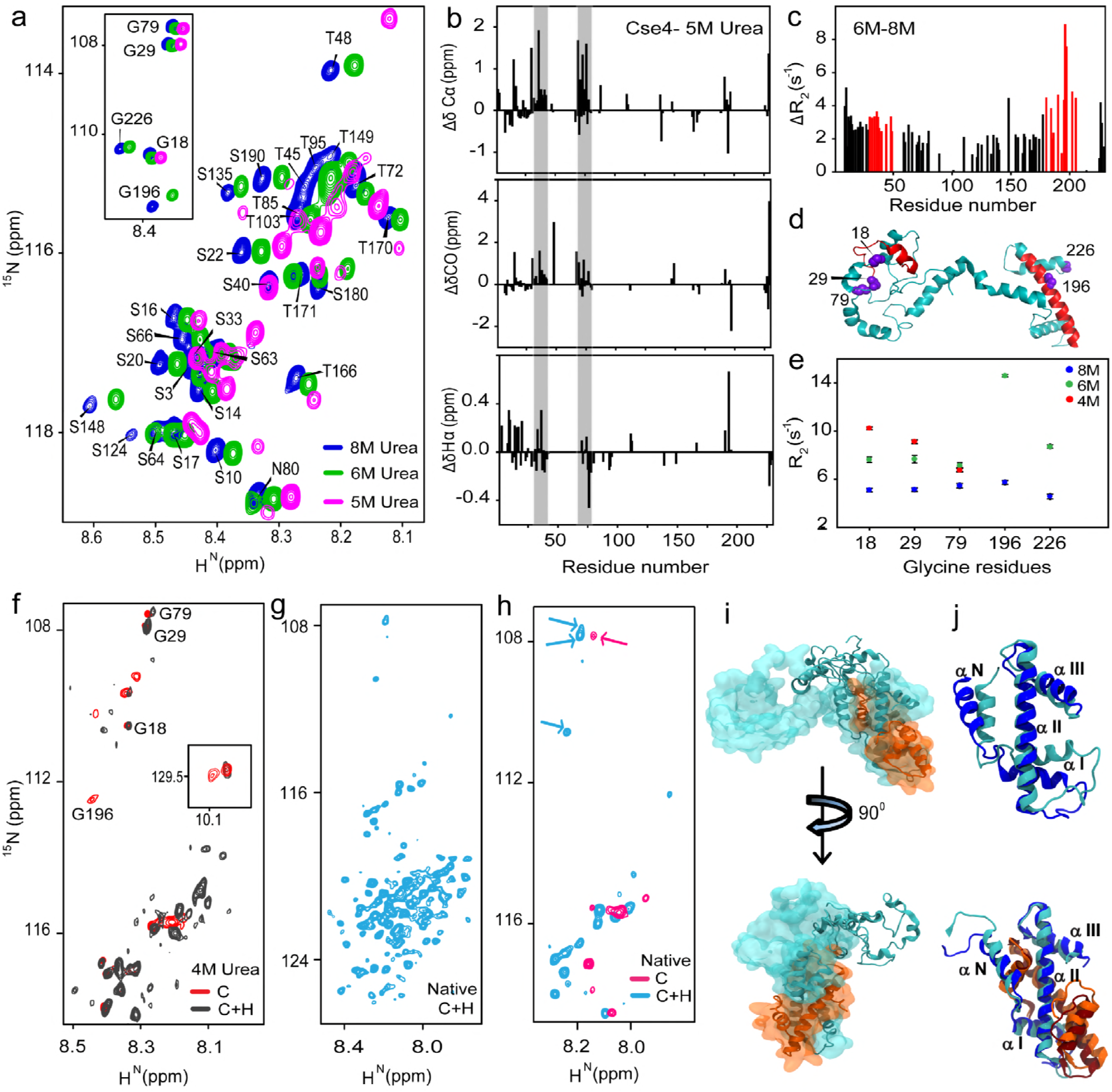
H4 stabilizes CENP-A^Cse4^. **a.** Representative regions of ^15^N-HSQC spectra of Cse4 at different urea concentrations, inset: peak positions of five Gly residues in similar conditions; **b.** Structural propensities of Cse4 in 5 M urea, plots of secondary chemical shifts from, ΔδCα, ΔδCO and ΔδHα, the residues D15-S18 (not marked), 30-42 and 75-78 show helical propensity (grey bars); **c.** Difference between the residue wise ^15^N transverse relaxation (R_2_) rates of Cse4 in 8 M and 6 M urea, residues showing maximum difference for each domain are marked in red; **d.** Modeled Cse4 structure[21] shows position of residues showing higher R_2_ difference (red), note that they are a part of the END and HFD regions, the Gly residues are marked as violet; **e.** ^15^N R_2_ for Gly residues in different denaturant concentrations; **f.** Overlapped regions of ^15^N-HSQC spectra of Cse4 and Cse4+H4 in 4M urea buffer, Gly residues are marked to show broadening of the C-terminus residues, inset: Trp side chain peaks; **g.** ^15^N-HSQC spectrum of Cse4+H4 complex without denaturant; **h.** Overlap between ^15^N-HSQC spectra of Cse4 and Cse4+H4 in the native state, the arrows indicate reappearance of the NTD Gly residues in the Cse4+H4 spectrum; **i.** Overlap of the structure of Cse4+H4 at the start (space filled) and end (ribbon) of simulation 1, the N-terminus does not fold back on C-terminus; **j.** Arrangement of the C-terminus helices with and without H4 binding at the start (Cse4-blue, H4 orange) and end (Cse4-cyan, H4-brown) of the simulation.

Next, we checked if the presence of H4 stabilized any of the domains of Cse4. Uniformly labeled ^15^N-Cse4 was co-folded with unlabeled H4, which resulted in increased the soluble fraction on removal of denaturant during folding. The (^15^N) Cse4+H4 complex was purified to yield the hetero-dimers adding up to ≈38 kDa, which is on the higher end of the detection limit of traditional solution-state NMR experiments; nevertheless important structural information about the two domains can still be acquired. At 4 M urea the resolution of Cse4 spectrum increased on co-folding with H4 (Fig 3f, Fig S6). The G196 disappeared and only one Trp side-chain peak was visible while the NTD Gly resonances superimposed with Cse4 monomer spectrum, suggesting that C-terminus is interacting with the H4. This trend continued with complete removal of denaturant, and the number of peaks increased in Cse4+H4 (0 M urea) in comparison to Cse4 monomer (Fig 3g, 2g). The NTD Gly resonances reappeared in the Cse4+H4 sample indicating a conformational change after H4 binding (Fig 3h). Cse4 C-terminus interacts with H4 [16], thus it is possible that CTD residues had broadened beyond detection and the NTD residues were reappearing because of the ‘release’ of N-terminus from the interaction with C-terminus when co-folded with H4.

The NTD does not interact with the CTD in any of the three MD simulations of Cse4+H4 complex (Fig 3i, Supplementary Movie 2, Fig S7a). The distance between the Trp residues is >4.8 nm in the presence of H4, indicating a lower probability of contact between the C and N terminus (Fig S7b), which is also evident by the contact maps (Fig S7c). The alpha-N and alpha-1 helix that were dislocated at the C-terminus in the monomeric Cse4 simulations, remained rigid in Cse4+H4 simulations with slight displacement in alpha-N (Fig 3j). The CTD of Cse4 was stabilzed by the presence of H4. This is critical not only for the nucleosome structure integrity but also for the various protein interactions where the Cse4 CTD is the binding interface. The structural rearrangement offers a plausible way of regulation at the centromere, where the CTD is oriented correctly and the NTD is ‘free’ to interact with other kinetochore proteins, only on H4 binding.

### H4 binding alters the conformation of the NTD

Co-folding of Cse4 and H4 indicated that the NTD no longer interacted with the CTD perhaps as a consequence of H4 binding at the C-terminus, as shown by the MD and NMR experiments. Next, we investigated if the binding of H4 to pre-folded Cse4 could alter the conformation of the NTD, which may be relevant in its interaction with other kinetochore proteins. We probed the change in conformation when H4 was added to native Cse4 to assess if it could ‘free’ the NTD from the CTD. There was a change in fluorescence lifetime of W178 at the CTD (Fig 4a, Table 3), but the change at the NTD was not significant, suggesting that H4 binds to the CTD and that the residues of the NTD were not involved. The flexibility of the CTD decreased slightly (Fig 4b,c), as W178 is restricted by the side chains of surrounding residues within Cse4 (Fig 2f) and H4 binding caused only a small change in the degree of its rotational flexibility (Fig S8). Surprisingly, the flexibility of the N-terminal Trp7 decreased significantly, as indicated by an increase in φ_1_ on H4 binding, (Fig 4b), signifying a local conformation change at the NTD on H4 binding. The lower β_1_ for the NTD in the presence of H4 also indicated reduced rotational freedom, which did not vary significantly for W178 (Fig 4c, Table 4). Addition of H4 dramatically increased the solvent accessibility of W7 (Fig 4d) proving that H4 could ‘free’ the NTD from interaction with the CTD. It should be noted that the dynamics of the Ala and Leu mutants differ slightly (Fig S9). The Ala mutant did not show much reduction in the amplitude of the short correlation time (Table 4) suggesting the presence of some transient interactions between the two domains. Histone H3 failed to induce a similar change in solvent accessibility of the NTD (Fig 4e) indicating that the effect is specific for H4. This observation has important consequences regarding the regulation of Cse4 targeting and functioning at the centromere. Only the specific partner can alter the conformation of the NTD such that it will be available for interaction with other proteins.

**Table 3:** Fluorescence intensity decay parameters of Cse4 mutants in presence of H4

| **Mutant** | **$\tau_1$ (ns) ($\alpha_1$)** | **$\tau_2$ (ns) ($\alpha_2$)** | **$\tau_3$ (ns) ($\alpha_3$)** | **$\tau_m$ (ns)** |
| --- | --- | --- | --- | --- |
| **W178L+H4** | 0.33(0.4) | 1.5(0.4) | 4(0.2) | 1.7±0.09 |
| **W7L+H4** | 0.46(0.4) | 1.8(0.4) | 4.4(0.2) | 1.8±0.04 |
| **W178A+H4** | 0.4(0.45) | 1.7(0.4) | 4.5(0.15) | 1.6±0.05 |
| **W7A+H4** | 0.5(0.5) | 2(0.36) | 4.6(0.14) | 1.6±0.07 |

**Table 4:** Fluorescence anisotropy decay parameters of Cse4 mutants in presence of H4

| **Mutant** | **$\phi_1$ (ns)** | **$\beta_1$** | **$\phi_2$ (ns)** | **$\beta_2$** |
| --- | --- | --- | --- | --- |
| **W178L+H4** | 0.33±0.01 | 0.42±0.06 | >20 | 0.58±0.06 |
| **W7L+H4** | 0.17±0.03 | 0.49±0.02 | 15±2 | 0.51±0.02 |
| **W178A+H4** | 0.3±0.06 | 0.45±0.08 | >20 | 0.55±0.08 |
| **W7A+H4** | 0.2±0.03 | 0.47±0.03 | 18.6±2.8 | 0.53±0.05 |

**Fig 4.**
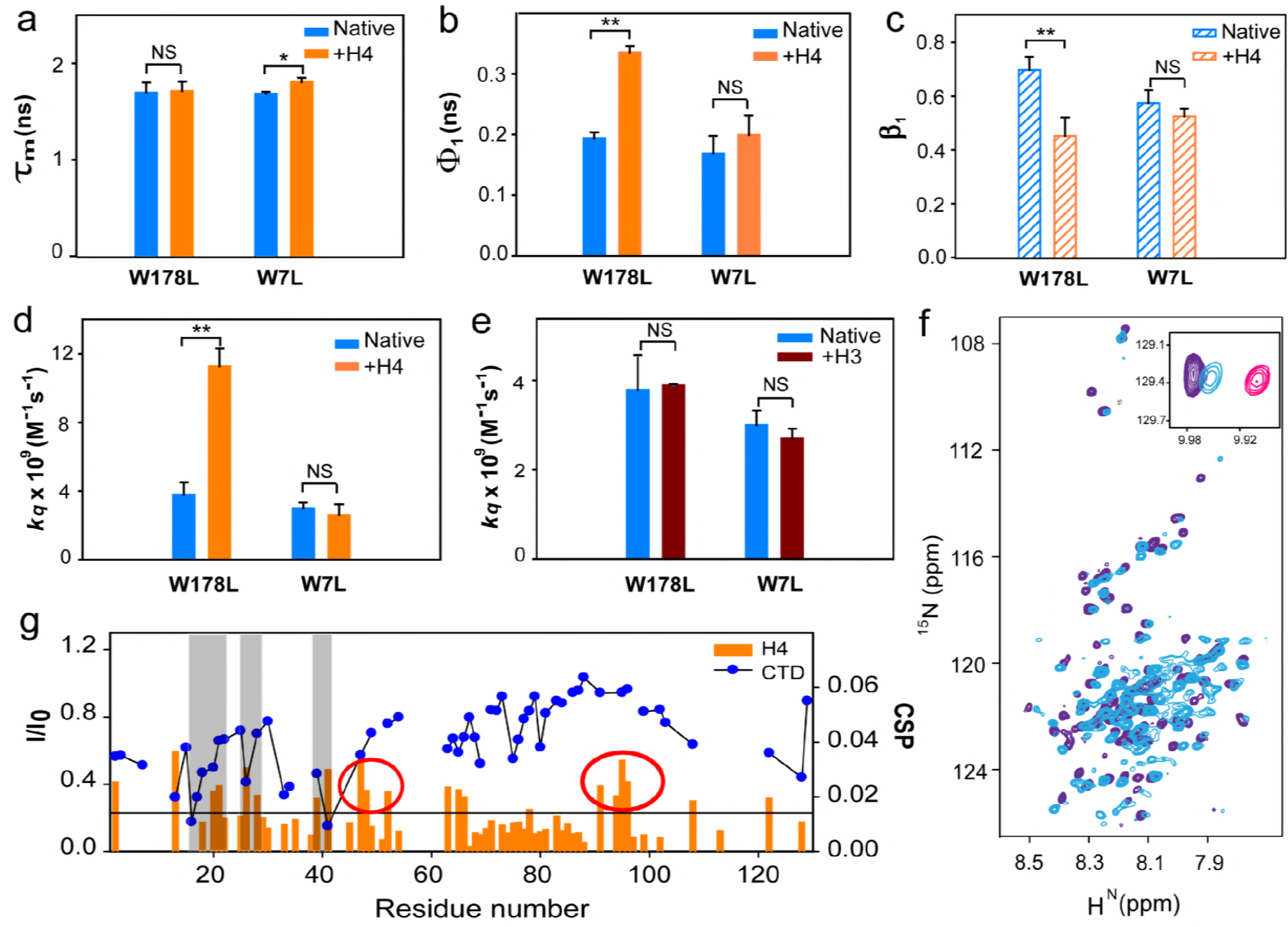
H4 binding alters NTD conformation. **a.** Comparison between the fluorescence lifetime values between Cse4 and Cse4+H4 for the two domains; **b.** Change in conformational flexibility of the Trp at the two domains with and without H4; **c.** β_1_ associated with the short correlation time for each mutant; **d**. Solvent accessibility of Trp residues at the two domains in Cse4 and Cse4+H4; **e**. Solvent accessibility of Trp residues at the two domains in Cse4 and Cse4+H3; **f**. Overlap of ^15^N-HSQC spectra of Cse4+H4 (light blue) and Cse4 (1-129) (purple), inset: Position of Trp side chain peak in Cse4, Cse4+H4 and Cse4ΔC. **g**. Comparison the residue specific CSPs calculated for NTD on addition of H4 (Orange histogram) and the change in intensity profile on titration with CTD (Blue scatter plot), grey bars represent the residues involved in interaction with both CTD and H4, some residues show significant CSP but do not interact with CTD (highlighted by red circles). Graphs-Blue, orange and brown represents Cse4, Cse4+H4 and Cse4+H3 respectively; mutants are specified at the X-axis. NMR spectra-Cse4+H4-light blue, Cse4-Pink, Cse4ΔC 1-129-purple. The statistical significance was calculated by one-way analysis of variance: *, p <0.05; **, p <0.01; NS (not significant), p >0.05; error bars represent SD.

Next, we recorded ^15^N-HSQC spectrum for truncated Cse4 NTD (Cse4ΔC, 1-129) to examine if the conformation of the NTD in full length Cse4 actually shifted towards a state similar to free NTD on addition of H4. Majority of the peaks were superimposable between the ^15^N-HSQC spectrum of Cse4+H4 and Cse4ΔC (res 130-229 deleted) (Fig 4f) confirming that the peaks that had broadened out in the full length Cse4 spectrum were from CTD and the NTD was free in the Cse4-H4 complex. The W7 side-chain peak for Cse4+H4 shifted closer to the Cse4ΔC than the Cse4 full length (Fig 4f, inset) showing that the NTD conformation was closer to its “open” state when H4 was added. Once the conformational plasticity was confirmed, we analyzed the residues involved in this interaction. Chemical shift perturbation (CSP) was observed for residues 20-28, 39-41, and 62-66 when Cse4ΔC was titrated against H4, indicating weak interaction between the two proteins. The residues from 94-129 also show CSP, but the resonances of all residues could not be included because of overlap in the spectrum. Interestingly, these residues that interact with H4 were also found to be a subset of those interacting with the CTD (7-51, 63-65, and 108-129), as seen by significant intensity change in the resonances that indicate intermediate to strong interaction (Fig 4g). Thus the NTD residues that are involved in a relatively strong binding with the CTD, also have a weak affinity towards H4 that may help in ‘opening’ the protein once H4 is added. This also explains the rigidity seen at the NTD on addition of H4. The residue wise interactions imply that the first step in the interaction between Cse4 and H4 would involve a weak affinity of H4 towards Cse4 NTD, which may help in dislodging the NTD from CTD before H4 stably binds to the C-terminus.

## Discussion

The role of Cse4 as an epigenetic marker for centromere identity is well established [26,27]. A conserved CENP-A Targeting Domain (CATD) consisting of loop 1 and helix 2 of the Histone fold domain in Cse4 is sufficient for maintaining centromere identity [14,15]. The N-terminal tail of Cse4 is dispensable for centromere targeting [28,29] but the deletion of the first 50 residues of Cse4 is lethal to cells [30]. A 33 residue stretch (28-60) that is required for interaction with other kinetochore proteins has been shown to be indispensible for cell survival [19] and its the role in the regulation of Cse4 levels in the cell is increasingly becoming more apparent [12]. The linear separation of the END from the HFD is not relevant, as the END fused directly to the CTD has been shown to confer wild type like functions [19]. However, some post translational modifications are known to occur at the NTD that regulate chromosome segregation and kinetochore integrity [31,32]. Thus, the actual significance of the length of the NTD in Cse4 and its structural organization is not clear.

We demonstrate here that in the soluble form, Cse4 exists in a ‘closed’ conformation, as a result of interaction between the NTD and the CTD. This interaction causes a change in the Histone-fold domain of the CTD where the alpha-N and alpha-I are dislocated. We believe that this change in the positions of helices of the CTD may interfere with Psh1 binding by changing the binding interface between the two proteins; since, Psh1 is known to interact with the CATD region of Cse4 and ubiquitinates four lysine residues in the CTD [9]. A peptidyl-prolyl cis-trans isomerase Fp3 is required to facilitate Psh1-mediaited degradation [33]. It is hypothesized that in the cellular environment the interconversion of the P134 of Cse4 by this isomerase from the ‘cis’ form to the ‘trans’ form ensures that any soluble Cse4 is targeted towards Psh1 mediated degradation, if not protected by the chaperone Scm3. Our data demonstrates a ‘closed’ conformation of Cse4 where the target residues for Psh1 (K131, K155, K163, and K172) might be inaccessible (similar to the ‘cis’ form) rendering Cse4 resistant to Psh1 mediated degradation. Our observations thus provide a rationale for the background levels of Cse4 observed in various cellular and biochemical assays. How this closed conformation subsequently interacts with the chaperone protein Scm3 will be interesting to study, given the fact that Scm3 deposits the dimer/tetramer onto DNA[34]. Crucially, in this conformation the NTD will not be free to interact with any kinetochore components and this could potentially negate the free monomeric Cse4 to nucleate kinetochore formation to any ectopic chromatin sites. Our data presents structural insights on retention of ‘inert’ soluble Cse4 molecule in the cell that is protected from proteolytic machinery and is also incapable to make any centromeric contacts. This indicates a novel mechanism of safeguarding Cse4 monomer, soon after its biogenesis, from proteolysis and mis-targeting (Fig 5).

**Fig 5.**
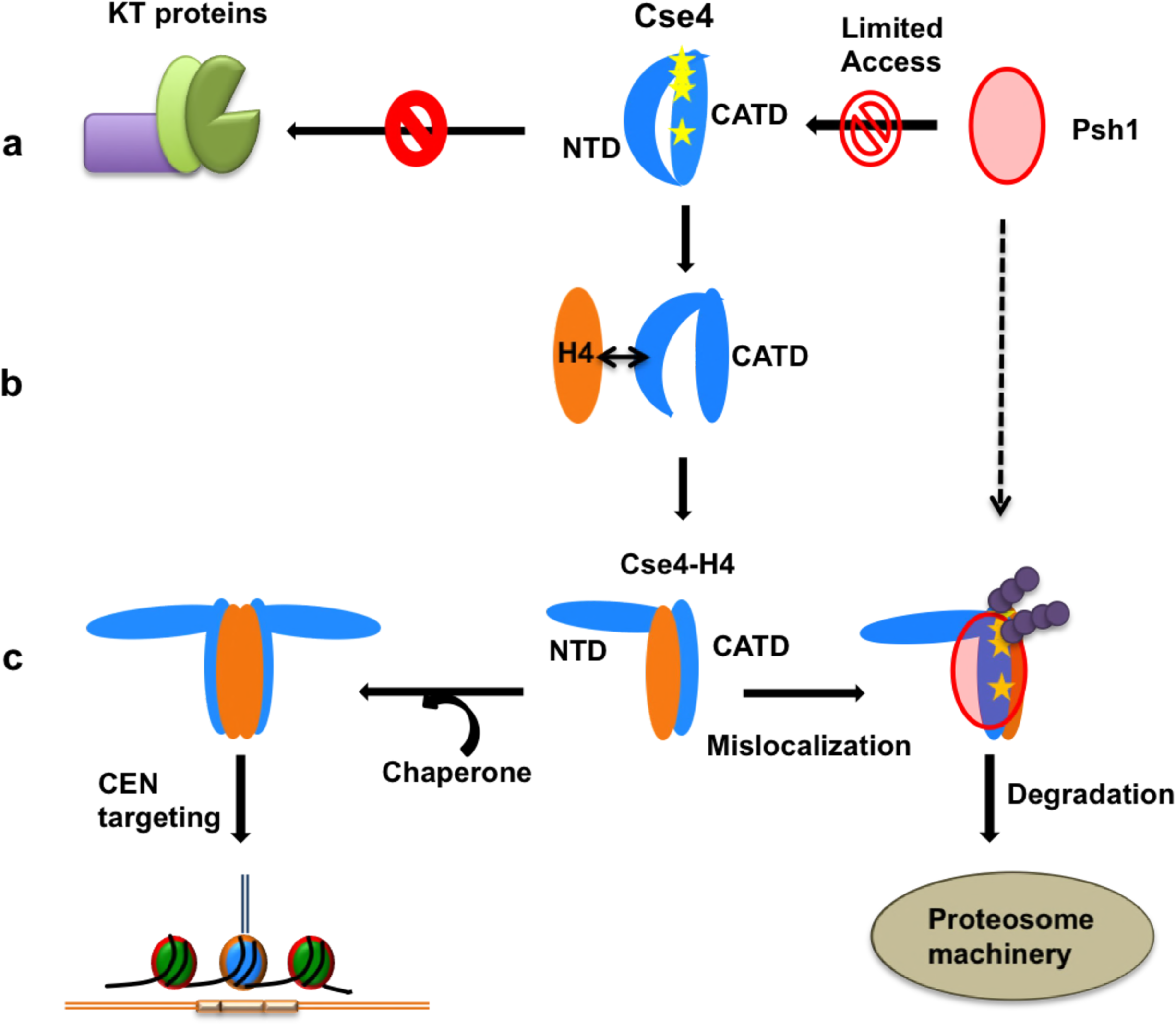
Proposed role of the NTD conformational change on regulation of Cse4. **a.** The closed state of Cse4 NTD prevents interaction with kinetochore proteins. It also alters the positions of the helices in the C-terminus thereby masking the binding sites (Lys residues highlighted as yellow stars) for Psh1. **b**. A possible transient state, when H4 comes in contact with Cse4. Affinity of NTD residues for H4 can help in dislodging the NTD from the CTD before H4 stably binds to the C-terminus; **c**. On H4 binding the NTD adopts an ‘open’ conformation. The Cse4-H4 dimer/tetramer can be deposited on the centromere by the chaperone Scm3, where the NTD is free to interact with the kinetochore. The C-terminus helices are re-oriented in a manner that facilitates ubiquitination and further degradation if Cse4-H4 mis-localizes to an ectopic location.

We observe that the conformational flexibility of both the domains of Cse4 is regulated by H4 binding. The positions of the alpha-N and alpha-I helices are retained close to their possible conformation in the nucleosome on H4 binding in our simulations. A recent study has shown that H4 facilitates the proteolysis of Cse4 by affecting its interaction with Psh1[35]. Our data are in agreement with this report, as the dissociation of the NTD from the CTD due to H4 binding would expose Psh1 binding sites and the complex would be targeted for degradation if localization to an ectopic site occurs. We report an allosteric effect on Cse4 upon H4 binding, as the interaction of H4 at the CTD causes a structural change at the NTD as shown by the fluorescence and NMR experiments. The NTD attains a ‘free’ conformation, which is evident by an increase in solvent accessibility and the overlap of ^15^N-H^N^ resonances of free NTD and Cse4-H4 complex spectrums. Crucially, histone H3, which is structurally similar to H4, does not change the NTD conformation, indicating that this interaction is specific to H4. The structural rearrangement observed here could be one of the mechanisms of regulation of specialized nucleosome formation at the right time. Only on H4 binding to ‘inert’ Cse4, the NTD is released for interaction with the kinetochore proteins or possibly even with DNA. This release will allow the NTD to interact with various proteins at different time points in response to cell cycle cues. Overall, this process prevents premature nucleation of kinetochore assembly in absence of H4, though further experiments are required to verify this. In conclusion, this study reveals that conformational flexibility of the NTD may act as a regulator for the correct localization of Cse4 in cells thereby preventing Cse4 from interacting with kinetochore components at ectopic locations.

## Materials and Methods

### Plasmids and Mutagenesis

Yeast *Saccharomyces cereviseae* Cse4 full-length protein cloned in pKS387 plasmid and Histone 4 and Histone 3 (*Saccharomyces cereviseae*) cloned in pET3a were kindly provided by Dr. K. Luger (University of Colorado Boulder). Single tryptophan mutants of Cse4 protein were created by site-directed mutagenesis using Kappa HiFi PCR kit (Kappa Biosystems, MA, USA). The mutants were selected by DpnI digestion (New England Biolabs). Plasmid DNA used for PCR was construct of pKS387 vector containing insert of full length Cse4. Tryptophan mutations were made at positions 7 and 178 (W7A, W7L, W178A, W178L). The truncated Cse4 construct Cse4ΔC (N-terminus 1-129) and Cse4ΔN (C-terminus 130-229) were cloned in pGEX6 vector. The clones have a GST tag with a precision cut site.

### Protein purification

The purification of H4, H3, Cse4, and its mutants was carried out according to Luger’s protocol [36]. Briefly, the cells were harvested and incubated with lysozyme at room temperature. The lysate was sonicated and centrifuged at 20,000 g for 20 min. The pellet obtained was washed twice with wash buffer (20 mM Tris pH 7.5, 100 mM NaCl, 1 mM EDTA, 1 mM PMSF) containing 1% triton X-100 and then washed with wash buffer (without Triton X-100). The remaining inclusion body pellet was dissolved in Guanidine hydrochloride (7 M). The sample was dialyzed against SAU-200 buffer (20 mM Sodium acetate pH 5.5, 8 M urea, 200 mM NaCl, 1 mM EDTA, 5 mM BME) and applied to the SP-Sepharose-fast flow column; any remaining impurities were removed by gel filtration (Sepharose-200). Protein purity was checked by SDS-PAGE. The proteins were dialyzed against distilled water with 10mM BME, the secondary structure of the proteins was checked by CD spectroscopy. For experiments with different urea concentrations, the proteins were dialyzed in acetate buffer containing required concentration of urea. The samples were maintained at pH 7 for all experiments and at pH 6.5 for NMR experiments.

The purification of GST tagged N-terminus (Cse4ΔC) and C-terminus (Cse4ΔN) was done using the Sepharose™ 4B system (GE Healthcare). The cell supernatant was and kept for binding with the beads for 3 h at 4°C in lysis buffer (20 mM phosphate pH 8, 100 mM NaCl, 1 mM EDTA). The beads were washed with wash buffers containing increasing amount of salt (100 mM, 300 mM, 500 mM NaCl). Precission enzyme was used to cleave the GST tag from the protein. The purity was checked by SDS-PAGE. Cse4ΔC was maintained at pH 5.5 for NMR experiments.

### Steady state fluorescence experiments

Fluorescence measurements were performed on a Perkin-Elmer spectrofluorometer, equipped with a data recorder and a temperature controlled cell holder. The fluorescence spectra were measured at a protein concentration of 20 µM with a 1 cm path length cell and at constant temperature (25°C). The samples were excited at 280 nm and emission spectra were recorded in the 290-500 nm range. The excitation and emmision slit width were set to 3 nm.

### CD measurements

Secondary structure of the different proteins was analyzed using CD. 20-30 µM of the proteins in the respective buffers was used. Far-UV circular dichroism (CD) spectra of the protein at 25°C were recorded on a JASCO-J-1500 CD spectrometer (Easton, Maryland). The samples were probed using 0.1 cm path-length quartz cell (Starna, Hainault, London), using a 1 nm bandwidth. For samples containing urea, scans were acquired from 210 nm to 260 nm, for all other samples 198-260 nm wavelength range was used. For signal averaging, three independent readings were taken. Raw data were processed by spectra smoothing and subtraction of respective buffers.

### Time resolved fluorescence measurements

The time-resolved fluorescence intensity as well as anisotropy decay experiments were carried out using a rhodamine 6G dye laser (Spectra Physics, Mountain View, CA), pumped by an Nd:YAG laser (Millenia X, Spectra Physics) and a time correlated single-photon counting (TCSPC) set up coupled to micro-channel plate photomultiplier (model R2809u; Hamamatsu Corp). Pulses (1ps duration) of 885 nm radiation from the rhodamine 6G dye laser were frequency tripled to 295 nm by using a frequency doubler (GWU, Spectra physics). The samples were excited at 295 nm and the emission was measured at the emission maxima (λmax) of the respective proteins, determined from their steady state fluorescence spectra. All the measurements were carried out on 30-50 µM samples. The instrument response function (IRF) was obtained at the wavelength 295 nm using a diluted colloidal solution of non-dairy coffee whitener. For time-resolved fluorescence intensity decay experiments, peak counts of 10,000 was collected with the emission polarizer oriented at the magic angle (54.7°) with respect to excitation polarizer. In time-resolved fluorescence anisotropy decay experiments, peak counts of 10,000 was collected with emission polarizer oriented at 0° (Parallel) and 90° (perpendicular) with respect to excitation polarizer.

### Calculation of the mean fluorescence lifetime

The fluorescence lifetime was analyzed by a method based on Levenberg-Marquardt algorithm [37]. The observed decay was deconvulated with the IRF to obtain the intensity decay function represented as a sum of three exponentials.

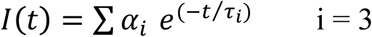

where I(*t*) is the fluorescence intensity collected with the emission polarizer oriented at magic angle (54.7°) at time *t* and α_*i*_ is the amplitude of the *i*^th^ lifetime τ_i_ such that Σ α_*i*_ = 1.

The mean fluorescence lifetime is calculated as

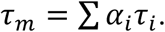

The goodness of fits were assessed from the reduced χ^2^ values and from the randomness of the residuals obtained from analysis.

### Fluorescence anisotropy decay kinetics

The anisotropy was calculated from experimentally obtained *I*_∥_ (*t*) and *I*_┴_ (*t*) using the equation:

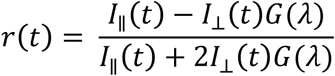

where r(t) is the time dependent anisotropy, *I*_∥_ (*t*) is the fluorescence intensity collected with emission polarizer at 0° (parallel) with respect to excitation polarizer, *I*_┴_ (*t*) is the fluorescence intensity collected with emission polarizer at 90° (perpendicular) with respect to excitation polarizer, and G(λ) is the geometry factor at the wavelength λ of emission. 50 µM solution of NATA (N-acetyl tryptophanamide) was used to calculate the G(λ) for the optics.

The *I*_∥_ (*t*) and *I*_┴_ (*t*) were fitted based on a model that assumes uniform motional dynamics in the sample, with each protein molecule associated with two rotational correlation times[23]:

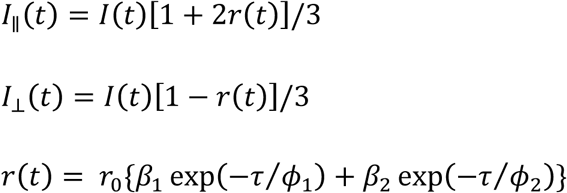

where r_0_ is the initial anisotropy i.e. in absence of any rotational diffusion (0.3) and β*i* is the amplitude associated with the i^*th*^ rotational correlation times ϕ_i_, such that Σ *β*_*i*_ = 1. The two correlation times can be interpreted to be associated with the local motion (short correlation time, ϕ_1_) and the global motion (long correlation time, ϕ_2_). The goodness of fit was assessed from the χ^2^ values.

### Acrylamide quenching of fluorescence

The protein concentration for quenching experiments was kept at 10 µM. The steady state fluorescence set-up was used for measurement. The protein aliquots were mixed with increasing concentration of acrylamide (0-0.3 M) and Trp fluorescence spectra were recorded for each sample. The maximum fluorescence intensity (F) was noted for each sample and the data was plotted according to the Stern-Volmer equation:

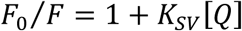

where F_0_ is the maximum fluorescence intensity without acrylamide, K_SV_ is the Stern-Volmer constant and [Q] is the concentration of the acrylamide in M. The bimolecular rate constant (*k*_*q*_) was calculated using the equation:

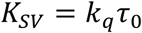

where τ_0_ is the mean life time value of the protein in absence of acrylamide.

### NMR experiments

Uniform ^15^N and/or ^13^C labeled samples were prepared by culturing the cells expressing Cse4 (full length and truncated (1-129)) protein in Minimal (M9) media supplemented with ^15^NH_4_Cl or ^15^NH_4_Cl and ^13^C labeled glucose. The purification was done as described above. For the protein folding studies, the Cse4 full-length sample was buffer exchanged to respective urea concentrations. The folded Cse4 full-length protein was prepared by dialyzing the sample against distilled water containing 2 M Arginine and 10 mM BME. D_2_O was mixed in 90:10 (H_2_O/D_2_O) ratios before recording spectra. pH of all Cse4 full length samples was maintained at 6.5 and Cse4 truncated sample (Cse4ΔC) was maintained at pH 5.5. The proton chemical shifts were referenced using DSS as an external calibration agent at 0.0 ppm whereas ^15^N and ^13^C were referenced indirectly as per BMRB protocol. The sample temperature was maintained at 25°C.

NMR experiments were recorded on Bruker Ascend 750MHz spectrometer. 5mm TXI probe with Z-gradient and deuterium decoupling was used to record a series of 2D and 3D experiments. 2D [^1^H, ^15^N]-HSQC was recorded for the full length Cse4 and truncated Cse4. The following experiments were recorded for assignment of Cse4 protein in 5M urea sample buffer-2D [^1^H, ^15^N]-HSQC, 3D HNCACB, HNCOCACB, HNCO, HNCACO, TOCSY-HSQC. The Cse4ΔC sample was maintained in 20 mM phosphate buffer with 150 mM NaCl and the following experiments were recorded for resonance assignment-2D [^1^H, ^15^N]-HSQC, 3D HNCA, HN(CO)CA, HNCO, HN(CA)CO, CBCA(CO)NH, TOCSY-HSQC, and H(CCO)NH. All spectra were processed with Topspin 2.1 version and analyzed with CCPNMR 2.3.1[38].

To analyze the secondary structure propensities of Cse4 protein in 6 M urea and 5 M urea the sequence corrected secondary chemical shifts (Δδ) for Hα, Cα and Cβ were calculated [39]. The random coil chemical shifts were taken from Schwarzinger et al. that uses 8 M urea at pH 2.3, 20°C for measurements on peptides to arrive at the random coil chemical shifts[40]. To investigate the backbone dynamics of Cse4, ^15^N relaxation data (R_2_) sets were recorded at 750 MHz frequency on a uniformly ^15^N labeled denatured Cse4 as well as Cse4 equilibrated with 6 M urea. ^15^N transverse relaxation rates (R_2_) were measured using delays, 10, 25*, 50, 90, 120, 150, 180*, 220 ms, where asterisk indicates duplicate measurements. Duplicate measurements for two random points were carried out for the verification of the error estimates. The cross peak intensities were measured as peak heights, using CCPNMR 2.3.1, which was also used to fit the relaxation data. The fitting was done to a single exponential decay function 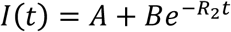 to extract the R_2_ values.

For the interaction studies, various sets of ^1^H-^15^N HSQC spectra were recorded at pH 5.5 with increasing equivalents of binding partner (CTD, and H4). The extent of interaction with each component was analyzed by checking the change in intensity profile and Chemical Shift Perturbation (CSP). The intensity profile of the amide cross-peaks affected during titration experiments was calculated by comparing their intensities (I) with those of the same cross-peaks (I_0_) without any addition, the data was normalized for dilution effect. The perturbation of amide cross-peaks chemical shifts during the interaction was calculated using the formula:

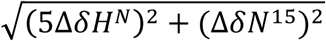

Residues showing CSPs greater than 2σ were considered significant[41].

### Molecular Dynamics

All the simulations were performed using GROMACS 4.6 package[42–45] with Bloom et al. as the source for the starting structures for monomeric Cse4 and Cse4+H4 simulations. Each protein system was placed in a dodecahedron box with distance between the protein and the box surface as 1nm. Amber99sb forcefield was used for the protein[46]. Simulation box was solvated using TIP3P water[47] and Na and Cl ions were added to achieve the salt concentration of 150 mM. The total number of atoms in Cse4 and Cse4+H4 systems were 92, 721, and, 118, 227 respectively. To enable the use of a 4 fs time step, all bond-angle hydrogens were treated using virtual sites[48]. Each protein system was energy minimized using steepest descent algorithm until the maximum force was less than 1000 kJ/mol/nm. Energy minimized structures were subjected to 100 ps temperature equilibration to 298 K using Berendsen thermostat[49] with a tau-t of 0.1 ps followed by pressure equilibration to 1 atm using Berendsen barostat[49] with tau-t of 1 ps. The final structure from pressure equilibration was used as the starting structure for production run simulations, where temperature was maintained using v-rescale thermostat[50] and Parrinello-Rahman barostat[51] with tau-t of 1 ps and tau-p of 5 ps. Four independent simulations for Cse4 and three simulations for Cse4+H4 from the corresponding pressure equilibrated structures were started with different starting velocities and from each simulation data was collected for 300 ns at 40 ps intervals. Analysis was done using tools from GROMACS package and in-house Python programs. Contact maps were generated using g_contacts tools using data from 250 ns-300 ns of each simulation[52].

## Acknowledgements

We thank Prof. K. Luger (University of Colorado Boulder, USA) for providing the plasmids for wild-type Cse4, H4 and H3. The authors acknowledge HF-NMR facility funded by Research Infrastructure Facility Committee, Industrial Research and Consultancy Center, IIT Bombay for NMR time, Dept. of Chemical Sciences, Tata institute of Fundamental Research for access to the Time correlated single-photon counting (TCSPC) set up, and CDAC PARAM YUVA II Supercomputing facility for computational time. The authors would like to thank Dr. Rohit Mittal (MRC, Laboratory of Molecular Biology, Cambridge, UK) for his critical feedback on the manuscript. NM is thankful for financial assistance from Council of Scientific and Industrial Research, India. AK is grateful for the Seed Grant provided by Indian Institute of Technology Bombay.

## Author Contributions

N.M. created Cse4 mutants and truncated proteins, conducted fluorescence and NMR experiments and analyzed the data. S.D. performed the molecular dynamics simulations and analysis. M.K. assisted in acquiring fluorescence lifetime data. G.K. assisted in acquiring and analysis of fluorescence data. S.K.G. helped in attaining clones and advised on biological analysis. N.M. and A.K. conceptualized the project and wrote the manuscript. N.M., S.D., G.K., S.K.G., and A.K. edited the paper. A.K. supervised the research.

## Conflict of interest

The authors declare no conflict of interests.

